# Ozone Stress Responsive Gene Database (OSRGD ver. 1.1): a literature curated database for crop improvement

**DOI:** 10.1101/2021.04.04.438353

**Authors:** Yadav Prachi, Usha Mina

**Affiliations:** School of Environmental Sciences, Jawaharlal Nehru University, New Delhi, India-110067

**Keywords:** Ozone stress, Bio-curation, Gene database, Ozone stress responsive genes (OSRGs)

## Abstract

Ozone (O_3_) is a major abiotic stress which severely affects the growth and development of plants. In order to cope up with ozone stress, plants exhibit a plethora of morphological, physiological and molecular changes. Various molecular studies have been performed and a variety of genes exhibiting expression in response to ozone stress have been identified. However, no existing database has been designed yet to present information on ozone stress responsive genes differentially expressed across different plant species. The lack of a curated database limits the research potential in the area and therefore a cohesive database should be designed as a data repository of the ozone responsive genes. OSRGD ver. 1.1 is a user friendly web interface to explore ozone stress specific transcriptome dataset of different plant species. It allows users to retrieve the ozone stress responsive gene data by keyword based search. Each entry upon keyword query contains detailed information about the specific species, its foliar injury symptoms, pattern of gene expression, and platform used for gene expression analysis with reference literature. This comprehensive biocuration will offer researchers a better biological insight into plants’ response to ozone stress which will focus on development of climate smart crops through crop breeding.

## Introduction

Tropospheric ozone (O_3_) is a highly oxidative secondary air pollutant produced through reactions between sunlight and primary air pollutants mainly nitric oxides, sulphur oxides, carbon oxides and volatile organic compounds (VOCs) emitted due to vehicular exhausts, fossil fuel burning and industries [Iriti and Faoro, 2008]. The effect of ozone on plants depends upon its exposure and uptake. Open stomata on the leaf surface serve as a major inlet for the ozone uptake [Tausz et al., 2007]. Only the amount of ozone reaching the apoplast or plasma membrane can have an effect on the tissue. After entry into the sub-stomatal cavity, ozone reacts quickly with cell wall and plasma membrane constituents and generates reactive oxygen species (ROS) burst in the apoplast [Rao and Davis, 2001; Kangasjärvi et al., 2005]. Ozone poses double jeopardy to plants in the form of low photosynthetic rates and impaired growth by long term exposure to low ozone doses and visible foliar lesions by short term exposure to higher ozone doses in ozone sensitive cultivars and species [Kangasjärvi et al., 1994].

Ozone being a potent oxidizing agent is capable of reacting with almost all bio-macromolecules, such as proteins, lipids and nucleic acids [Mustafa, 1990]. In plasma membrane, fatty acids are the main target for ozone which results in lipid peroxidation causing severe membrane damage by altering the membrane fluidity [Iriti and Faoro, 2008]. Ozone also inactivates the membrane ATPases and disturbs the overall permeability of the plasma membrane [Dominy and Heath, 1985]. Ozone results into oxidation of primarily tyrosine, tryptophan, methionine, cysteine and histidine amino acid residues leading to modification in structure and activity of proteins generating hydrogen peroxide and hydroperoxides [Freeman and Mudd, 1981]. Ozone is also responsible for the nucleic acid damage by oxidation of deoxyribose, strand breaks and modification of bases. DNA damage due to ozone was demonstrated by the stimulation of poly (ADP-ribose) synthetase, a chromatin-bound enzyme activated in response of DNA strand break [Hussain et al., 1985]. These oxidation products mediate reactions within the cells leading to identified ozone effects such as decrease in photosynthetic rates, increased electrolyte leakage and accelerated senescence and cell death.

Since oxidative stress is an unavoidable event, plants have developed interplay of protective antioxidative defense systems for the detoxification of these reactive oxygen species (ROS). The antioxidative defense systems include non-enzymatic antioxidants such as ascorbic acid [Smirnoff and Wheeler, 2000] and glutathione [De Kok and Tausz, 2001] and antioxidant enzymes such as superoxide dismutase (SOD, EC 1.15.1.1), catalase (CAT, EC 1.11.1.6) and glutathione peroxidase (GPX, EC 1.11.1.9) for the protection of cell from ROS induced damage [Ludwikow and Sadowski, 2008]. The ascorbate-glutathione cycle (AsA-GSH) plays a major role in combating the oxidative stress and form the first line of defense against ozone. The AsA-GSH cycle involves oxidation and reduction of ascorbate, glutathione, and NADPH catalyzed by ascorbate peroxidase (APX), dehydroascorbate reductase (DHAR) and glutathione reductase (GR). Ascorbate acts as an antioxidant buffer in the apoplast and can result in detoxification of up to 30–50% ozone uptaken by the leaves [Polle et al., 1995; Turcsányi et al., 2000]. The enzymatic components of the antioxidant systems are distributed majorly in different subcellular components, such as mitochondria, chloroplasts, cytosol and peroxisomes [Mittler et al., 2002]. These enzymes catalyze the ROS degradation and form the frontline in scavenging the toxic ROS levels.

Ozone stress causes foliar injury symptoms in the form of small, interveinal bronze, pale yellow or reddish stippling on the adaxial leaf surface, which extends to form necrotic spots, lesions and flecks [Günthardt-Goerg and Vollenweider, 2007]. The generation of visible leaf injury is a result of ozone-induced ROS accumulation in the cells which elicit a hypersensitive response (HR) resulting in lesion propagation and leaf necrosis [Iriti and Faoro, 2008]. Also the ozone-derived ROS act as signal molecules that activate the programmed cell death (PCD) pathway [Overmyer et al., 2005] (Fig. 1).

**Fig. 1:**
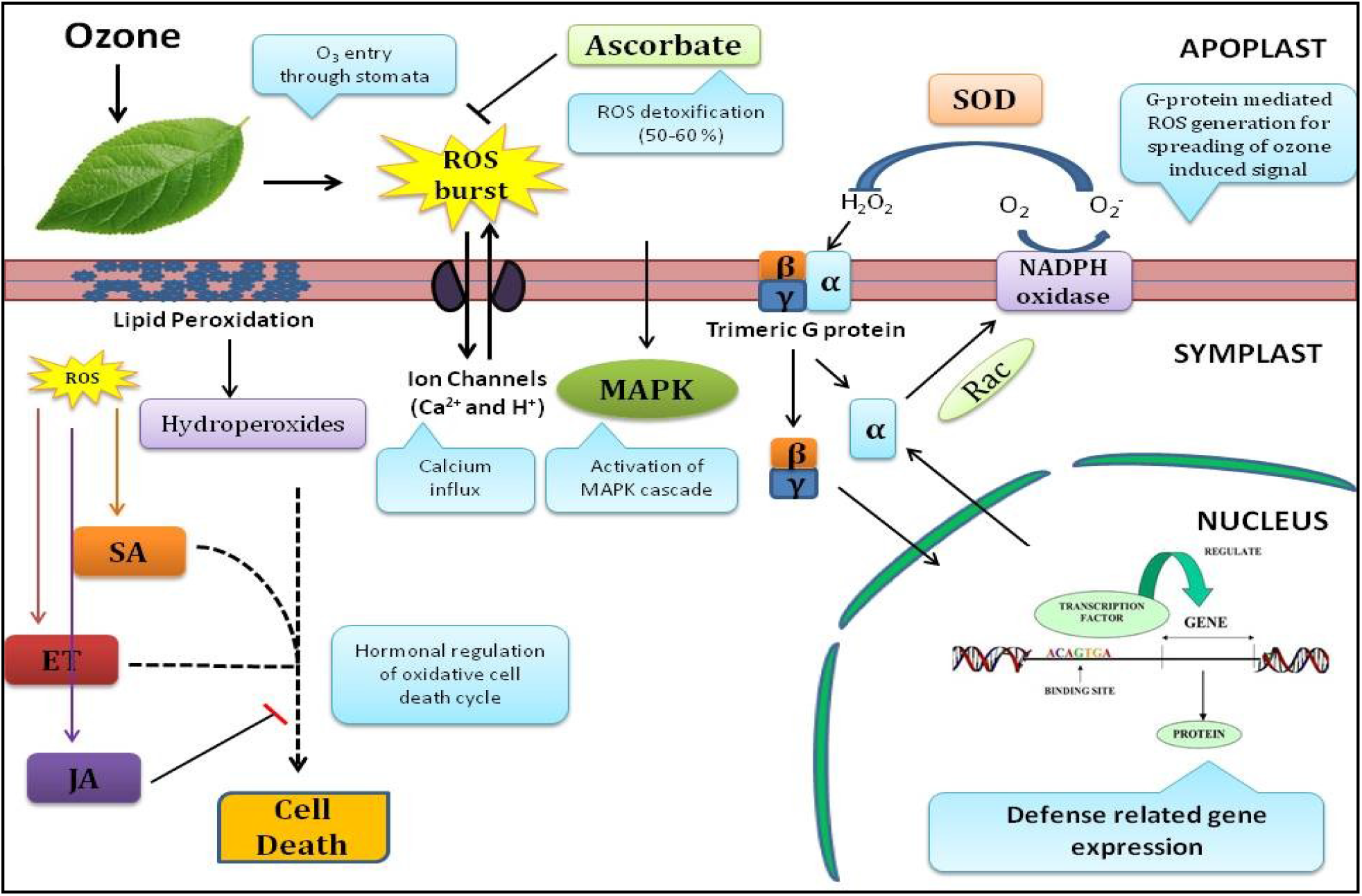
Ozone stress induced signaling pathways

Phytohormones viz. ethylene, salicylic acid, jasmonate and abscisic acid play a significant role in determining the initiation, propagation and containment of foliar injury [Overmyer et al., 2003]. There have been many reports indicating that ozone-induced foliar injury is a result of self amplifying oxidative cell death cycle driven by these phytohormones [Kangasjärvi et al., 2005]. Ethylene directs the endogenous ROS production which induces the salicylic acid dependent PCD pathway and lesion propagation [Ciardi et al., 2000]. The cell death continues until the jasmonate dependent pathway which is antagonistic to lesion propagation, confines the lesion enlargement [Berger, 2002]. The molecular mechanisms underlying ozone phytotoxicity have only started to get explored in recent years. The study of gene expression pattern underlying the ozone stress which governs the ozone tolerance mechanism has remained a key challenge for the researchers. For this the whole annotation of transcriptome is essential for quantifying the changes in expression level of each gene under the ozone stress.

During the past decade, a significant progress in plant genomics provided knowledge about the molecular mechanisms evolved by plants to survive the abiotic stress. The major genomic approaches to quantify the gene expression at transcriptome level include hybridization or sequencing based approaches. Hybridization based approaches involve custom made microarrays or commercial high density oligo microarrays. These microarray techniques are high throughput and relatively inexpensive but has the limitation of high background signals due to cross hybridization and reliance to the existing knowledge of the genome sequence. To overcome these limitations, the novel high-throughput DNA sequencing based approaches such as 454 pyrosequencing and RNA-seq have been developed as a revolutionary tool for transcriptome profiling [Wang et al., 2009; Marguerat and Bähler, 2010]. These approaches offer single base resolution with low background signal and provide digital gene expression levels at a much lower cost [Mutz et al., 2013].

Several studies have utilized these approaches for differential gene expression analysis in different plant species under ozone stress. In *Arabidopsis thaliana*, microarray analysis and RNA-seq were used to reveal the candidate genes regulating the response to ozone stress [Mahalingam et al, 2006; Xu et al., 2015]. The ozone responsive transcriptome of rice seedling and seed coat have also been studied using DNA microarray approach [Cho et al., 2008; Cho et al., 2013]. Illumina sequencing has been utilized to analyze the transcriptomic response to elevated ozone in legume crops viz. *Pisum sativum, Glycine max* and *Phaseolus vulgaris* [Yendrek et al., 2015]. *Glycine max* has also been extensively studied for transcriptome annotation under ozone stress for leaf tissue, seed coat, flower and pod [Leisner et al., 2014; Whaley et al., 2015; Leisner et al., 2017; Waldeck et al., 2017]. Transcriptome studies have also been done on *Medicago truncatula* and *Pak choi* using DNA microarray and RNA-seq respectively [Iyer et al., 2013; Zhang et al., 2017]. Apart from these crop plants, studies have also been performed for tree species such as *Populus* [Gupta et al., 2005; Street et al., 2011], *Quercus* [Natali et al., 2018; Soltani et al., 2020], *Betula pendula* [KONTUNEN-SOPPELA et al., 2010] and *Viburnum lantana* [Gottardini et al., 2016]. These studies have enriched the knowledge about genes which were differentially expressed under elevated ozone stress.

When a large number of genes across different plant species are associated with expression of an attribute, bio-curation can be an approach to identify important candidate genes from already existing information available in the public domain which can be exploited for crop improvement through breeding. There are few databases that are designed for stress responsive genes in plants. These include Arabidopsis stress responsive gene database [Borkotoky et al., 2013], PSRN [Li et al., 2018], PSPDB [Kumar et al., 2014], PASmiR [Zhang et al., 2013], PRGdb [Sanseverino et al., 2010], DroughtDB [Alter et al., 2015], QlicRice [Smita et al., 2011], RiceSRTFDB [Priya and Jain, 2013], PhytAMP [Hammami et al., 2009], STIFDB2 [Naika et al., 2013], etc. However no existing database has been designed to provide information on ozone stress specific genes which are differentially expressed across the various plant species under ozone stress.

In this study, we present the ozone stress responsive gene database (OSRGD ver. 1.1) which is first comprehensive manually curated database that can serve as a data source and tool to the scientific community to gather information about ozone stress responsive genes in various plant species. This biocuration aims to serve as a primary archive to efficiently manage ozone stress specific gene data and provide a platform for research community to explore data in more accessible, findable and usable form. The web interface is freely accessible at http://www.osrgd.com.

## Materials and Methods

### a) Data collection and compilation

The manual curation approach was used to extensively search the literature published in last 50 years (1970-2020) which is available on the public data sites such as Pubmed, Google Scholar, Refseek, Microsoft Academic, and Worldwidescience. After preliminary screening of the literature related to plants under ozone stress, the detailed information was further extracted by using different combinations of keywords (Table 1) as search conditions and web crawler bot was used to collect the abstracts of related literatures. Upon curation, a total of 40 literature studies were identified for 16 plant species in which complete gene expression profiling in different plant tissues and cultivars/varieties have been carried out till date. The relevant literature was downloaded and catalogue to acquire the required information. The collected literature served as a reference dataset for the study. Only those studies were included in the database which offers the complete dataset of differentially expressed genes under ozone stress. There were few studies where the gene dataset of ozone responsive genes was not publically available and therefore those studies were excluded from the web interface.

**Table 1:** Keyword combinations used in different search engines with search results

| S.No | Keywords used | Google Scholar | Pubmed | Refseek | Microsoft Academic | World wide science |
| --- | --- | --- | --- | --- | --- | --- |
| 1. | Plants and ozone stress | 1,01,000 | 6841 | 4,23,000 | 40 | 1676 |
| 2. | Plants under ozone stress | 76,600 | 6359 | 3,14,000 | 40 | 1375 |
| 3. | Gene expression in plants under ozone stress | 20,800 | 4090 | 1,17,000 | 2058 | 1105 |
| 4. | Gene regulation study in plants under ozone stress | 19,500 | 3689 | 1,48,000 | 1,029 | 1077 |
| 5. | Genes regulated under ozone stress in plants | 19,500 | 3016 | 1,76,000 | 37 | 977 |
| 6. | Ozone stress responsive genes in plants | 19,200 | 2152 | 1,62,000 | 12 | 1066 |
| 7. | Plant transcriptomics under ozone stress | 12,400 | 446 | 20,200 | 61 | 695 |
| 8. | Microarray under ozone stress in plants | 10,400 | 1123 | 20,600 | 29 | 636 |

The plant species identified in the literature included *Arabidopsis thaliana*, *Oryza sativa* (Rice), *Medicago truncatula* (Barrel clover), *Phaseolus vulgaris* (Common bean), *Glycine max* (Soybean), *Pisum sativum* (Garden pea), *Brassica Campestris* (Pak choi), *Capsicum annum* (Pepper), *Salvia officinalis* (Common sage), *Viburnum lantana* (Wayfaring tree), *Malus* (Crabapple), *Populus* (Poplar), *Betula* (Birch), *Quercus* (Oak), *Fagus sylvatica* (European beech), *Fraxinus pennnsylvanica* (Green ash) (Table 2).

**Table 2:** Plant species evaluated for ozone stress responsive gene database (OSRGD ver 1.1)

| S.No . | Species name | Common name | Category/ Growth habit | Tissues used in study | Ecotype/Cultivar used in study | Ecological significance | Reference |
| --- | --- | --- | --- | --- | --- | --- | --- |
| 1. | *Arabidopsis thaliana* | Arabidopsis | Herb | Leaf | Wassilewskija (Ws-0), C24 (Ozone tolerant), Te (Ozone sensitive), Columbia (Col-0), RLD1, CT101, Cape Verde Islands (Cvi-0), Landsberg erecta (Ler) and a relative, *Thellungiella halophila* (Th) | Model system in plant molecular biology and genetics | Li et al., 2006; Mahalingam et al., 2006; Tosti et al., 2006; Yoshida et al., 2009; Short et al., 2012; Xu et al., 2015 |
| 2 | *Oryza sativa* | Asian rice | Herb | Leaf, Seed, Panicle* | cv. Nipponbare, Koshihikari, Kirara397, Takanari, Kasalath, | Staple food of South Asia (Cereal crop) | Cho et al., 2008; Cho et al., 2013 |
| 3 | *Medicago truncatula* | Barrel clover | Herb | Leaf | cv. Jemalong (Ozone sensitive), JE154 (Ozone resistant) | Model organism for legume biology | Iyer et al., 2013 |
| 4 | *Pisum sativum* | Garden pea | Climber | Leaf | *Pisum sativum* L. | Vegetable crop | Yendrek et al., 2015 |
| 5 | *Phaseolus vulgaris* | Common bean | Herb | Leaf | *Phaseolus vulgaris* L. | Legume crop | Yendrek et al. 2015 |
| 6 | *Glycine max* | Soybean | Herb | Leaf, Flower, Pod, Seed coat | *Glycine max* L., *Glycine soja*, Jinpumkong, Cheongjakong, Mandarin (Ottawa), Fiskeby III, Pioneer 93B15 | Legume crop | Leisner et al., 2014; Whaley et al., 2015; Yendrek et al. 2015; Leisner et al., 2017; Waldeck et al., 2017 |
| 7 | *Brassica campestris* | Mustard | Herb | Leaf | cv. Jingguan (Pak choi) | Oilseed crop | Zhang et al., 2017 |
| 8 | *Capsicum annum* * | Pepper | Herb | Leaf | cv. Dabotop (ozone-sensitive) and cv. Buchon (ozone-tolerant) | Vegetable crop | Lee et al., 2006 |
| 10 | *Viburnum lantana* | Wayfaring tree | Shrub | Leaf | *Viburnum lantana* L. | Deciduous shrub | Gottardini et al., 2016 |
| 11 | *Malus* * | Crabapple | Tree | Leaf | Hongjiu | Deciduous | Wu et al., 2020 |

|  |  |  |  |  |  | fruit tree |  |
| --- | --- | --- | --- | --- | --- | --- | --- |
| 12 | *Populus* | Poplar | Tree | Leaf | *Populus deltoides* clone '546',<br>*Populus tremuloides* clone '216', F2 mapping population formed from cross between *P. trichocarpa* and *P. deltoides* | Deciduous flowering tree | Gupta et al., 2005;<br>Street et al., 2011 |
| 13 | *Betula* | Birch | Tree | Leaf | *Betula pendula* Clone 4 (Ozone sensitive)<br>Clone 80 (Ozone tolerant),<br>*Betula papyrifera* * | Deciduous hardwood tree | KONTUNEN-SOP<br>PELA et al., 2010 |
| 14 | *Fagus sylvatica* * | European beech | Tree | Leaf | Beech saplings | Deciduous tree (Ornamental) | Olbrich et al., 2005 |
| 15 | *Fraxinus pennsylvanica* | Green ash | Tree | Leaf | *Fraxinus pennsylvanica* L. | Deciduous tree | Lane et al., 2016 |
| 16 | *Quercus* | Oak | Tree | Leaf | *Quercus ilex* L.<br>*Quercus rubra* L. | Deciduous tree | Natali et al., 2018;<br>Soltani et al., 2020 |

\*The ozone responsive gene data was not publically available due to which these species were not included in the database.

Data related to these plant species such as family name, cultivar, ozone concentration and exposure period, foliar ozone injury symptoms, sampling time, platform used for gene expression study, data repository *Id* and list of genes up-regulated and down-regulated under ozone stress were extracted from available literature. The data was manually inserted into the database. The organizational structure of OSRGD ver. 1.1 has been illustrated in Fig 2.

**Fig. 2:**
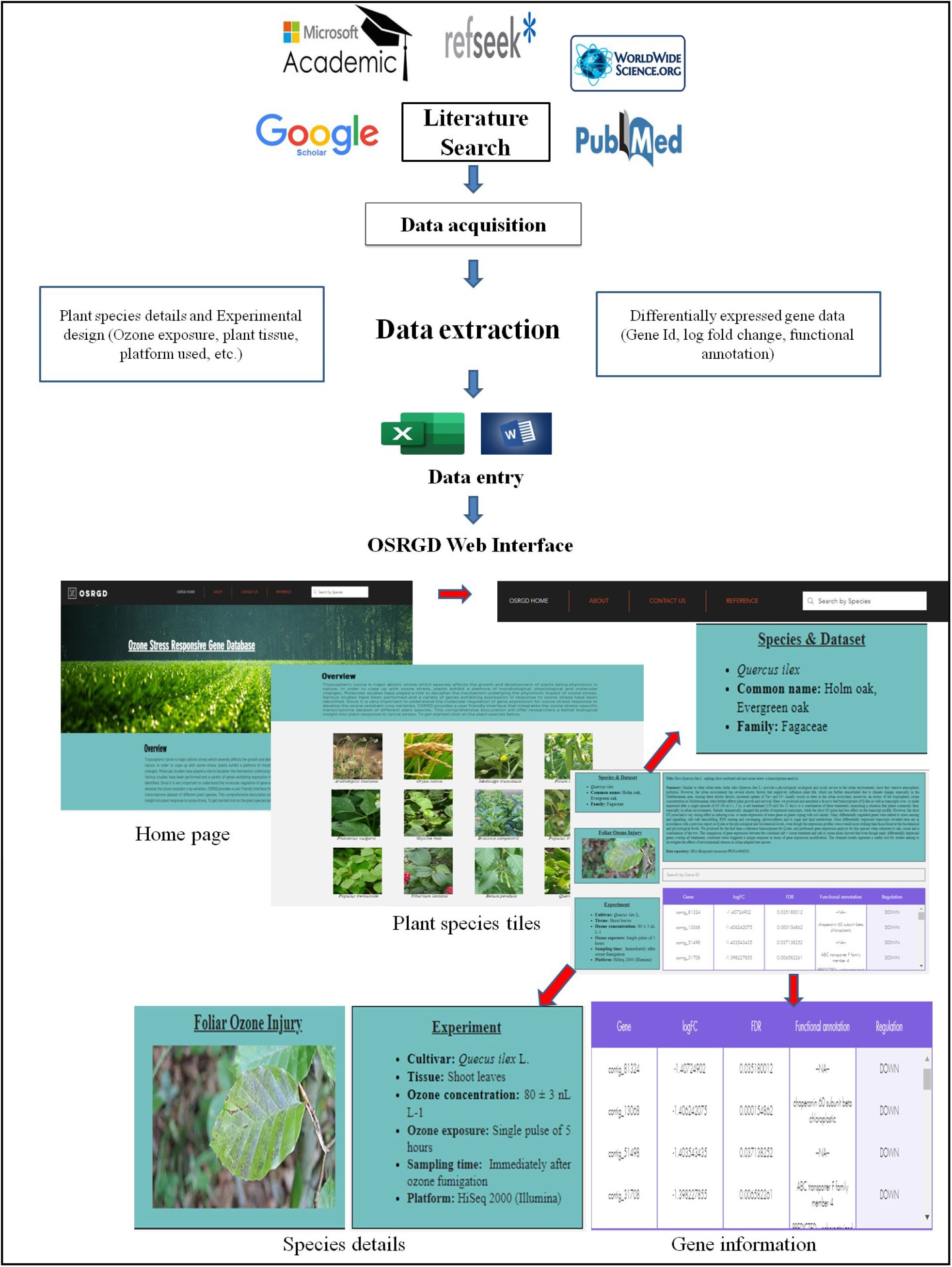
Organizational structure of OSRGD ver. 1.1

### b). Database web interface

The ozone stress responsive gene database (OSRGD ver. 1.1) is designed and developed in a user-friendly way using a comprehensive platform WIX which offers drag and drop tools for development of web interface. The database is designed in graphical user interface manner and is a structured representation of genes up-regulated or down-regulated in response to the ozone stress in different plant species. The user can access the web interface over the secure channel of internet via uniform resource locator http://www.osrgd.com. The front end interface is developed with the help of scripting languages such as html and JavaScript which are generally featured with speed, simplicity, popularity and inter-operability. The current web interface tends to be very fast and easy to use as it runs with in the user’s browser without the requirement of any outside resources. Currently all major browsers support just in time compilation for high level computational languages. Hence there is no need to compile the code of interface before executing it. Further the database is designed with the help of MS Excel which provides the structured view to the user. The tabular representation of the gene data increases the data visualization and provides a summarized view to the user. The web interface is designed in such a manner that it supports simplicity, openness and extensibility. Also the web interface was made mobile phone friendly for its easy access from a variety of digital devices.

### c). Database architecture

The extensive literature search and data mining resulted in collection of specific genes reported to be up- and down-regulated under ozone stress. The database can be queried using keywords for plant species name and Gene Id in the search box. Also plant species name along with their images are used as tiles on the home page for the easy access to the gene data. For each plant species, a separate page was created which consists of all the information related to the study and gene data in the form of spreadsheet.

The information in the database in divided into different sections of the web interface:

- **Homepage**: In order to keep the database interface simple and user-friendly, a simple menu was developed on the homepage with basic features. A brief overview of the database is given at the homepage to provide a short description about the database. A navigation menu is located (About, Contact us, References, Search box) at the top of interface to help visitors to track the information they are looking for. A chat box feature was also embedded into the web interface to enable the users to post their queries.
- **Species page**: The species which were evaluated in the database for ozone stress responsive genes were presented in the form of picture tiles for easy accessibility to the user. User can directly click upon the species tile to navigate on the page carrying species details. This feature omits the need to go to search menu every time and made the switching easy between different species.
- **Species details:** The page includes the information of the plant species with its scientific name, common name and family name to which the plant belongs to along with their foliar ozone injury symptoms under ozone stress. The experimental details described the cultivar and the tissue used for study along with the ozone concentration and exposure period of ozone stress. Also the platform used for gene expression studies under ozone stress was specified. For published gene data, the SRA data uploaded on NCBI (National centre for Biotechnology Information) or EBI (European Bioinformatics Institute) have been linked in OSRGD ver. 1.1.
- **Gene information:** Gene Id, Accession number, AGI code, Gene bank Id were given according to the platform used for the study. The gene up-regulation or down-regulation was depicted with the help of log fold change values (along p-value) where fold change value >2 and log2 fold change value >1 is interpreted as up-regulated and fold change value < 0.5 and log2 fold change value <1 is interpreted as down-regulated. Also the functional annotation for each gene was given to determine the functional role of the differentially expressed genes under ozone stress.

The architecture of this database also contains the facilitation web pages such as About, Contact Us, References and Tools. The About page addressed the basic questions associated with the ozone stress and the answers to frequently asked questions (FAQs) by the users. The Contact Us page helps the visitor to find the quick information about the authors. The list of literature references provided by clicking ‘References’. OSRGD ver. 1.1 also provides set of tools to facilitate further analysis of available gene data and to enable access to additional relevant information about the gene. In order to keep the web interface simple and light, we have not included the extensive structural images and graphs thus increasing its usability on various digital devices.

## Discussion

The extensive literature search resulted into compilation of genes differentially expressed under ozone stress in different plant species. Maximum numbers of studies were reported for *Arabidopsis thaliana* (model plant) to elucidate the genes involved in ozone stress induced oxidative signaling, Ca^2+^ signatures and phytohormone synthesis in different ecotypes. *A.thaliana* serve as an important model system for investigating the plant stress response which is attributed to its small genome size (~ 135 Mb), short life cycle and ease of generating mutants and transgenic plants. A variety of mutants and ecotypes are available for *A. thaliana* which adds to its utility in studies of plant response to ozone stress.

The temporal expression study in ozone sensitive *A.thaliana* ecotype Wassilewskija (Ws-0) showed biphasic ROS burst with a short peak at 4h and a higher peak at 16h. The whole genome expression profiling identified 371 differentially expressed genes under ozone stress. There was an early induction of proteolysis and hormone responsive genes indicating the rapid activation of oxidative cell death pathway which suggests a significant role of phytohormones in ozone induced cell death. There was also a down-regulation in genes involved in carbon utilization, metabolic pathways and signaling indicating an ineffective defense response by the Ws-0 ecotype [Mahalingam et al., 2006]. Another study showed the up-regulation of several WRKY genes in *A.thaliana* ecotype Col-0 which strongly indicated that WRKY transcription factors are key factors of signal transduction in ozone treated plants [Tosti et al., 2006]. Another study with Cvi-0, Ws-0 and Col-0 ecotypes of *A.thaliana* demonstrated development of foliar lesions in Ws-0 and Col-0 within 10 days of ozone treatment. However, Cvi-0 showed up-regulation of chaperons, receptor kinase-like and photosynthesis related genes along with AtSR (a mammalian NFκB homologue), WRKY and C2H2 protein and antioxidants. Whereas Ws-0 exhibited down-regulation of two AtSR, C2 domain proteins and genes involved in the growth of cell wall [Li et al., 2006].

Yoshida et al. showed that de novo biosynthesis of glutathione is one of the significant defense responses during ozone stress which is largely controlled by salicylic acid and ethylene [Yoshida e al., 2009]. A stimulus specific calcium signature regulating the transcriptional response to ozone stress was also identified in Col-0 ecotype of *A.thaliana* [Short et al., 2012]. An antagonistic relationship between salicylic acid and apopalstic ROS signaling was identified regulating the gene expression in response to ozone stress. It was shown that plants are able to prioritize the ozone response between salicylic acid and ROS signaling based on the exposure duration and dose of ozone. Another study by Xu et al. identified the role of defense hormones in regulation of gene expression network and cell death under ozone stress by using ozone sensitive and tolerant *A.thaliana* ecotypes Te and C24, respectively. It was shown that in response of apoplastic ROS, no single pathway is responsible for gene expression regulation. Instead, a combination or overlapping of several different pathways regulates the ozone stress induced gene expression and cell death [Xu et al., 2015].

The gene data from different ecotypes of *A.thaliana* was subjected to web tool Venn (http://bioinformatics.psb.ugent.be/webtools/Venn/) to study which genes are in intersection or are unique to a particular ecotype (Fig. 4). The up-regulated genes were identified from the list of unique genes of each ecotype using VLOOKUP function in MS Excel and the identified genes were then subjected to gene ontology classification using GO term enrichment at TAIR (https://www.arabidopsis.org/tools/go_term_enrichment.jsp). Very few genes were found to be in intersection suggesting that different *A.thaliana* ecotypes have distinct gene regulation pattern under ozone stress. The Ws-0 ecotype exhibited the maximum up-regulation in genes responsible for defense response to bacterium indicating the resemblance of ozone induced response to pathogen stress [Kangasjärvi et al., 1994]. In Col-0 ecotype, the gene expression was up-regulated for camalexin metabolite biosynthesis contributing to the resistance of *A.thaliana* against bacterial and fungal pathogens. In ecotype C24, the most enriched biological process is pattern recognition receptor signaling pathway upregulated under innate immunity mechanism of plants to deal with pathogens. Whereas, ecotype Te had the most up-regulated genes involved in proteolysis by ubiquitin-proteasome system. The ubiquitine-proteasoe system has a major role in plants response and adaptation to abiotic stress by influencing the production and signaling of stress related hormones [Stone et al., 2019]. The study of different Arabidopsis ecotypes suggested that plant response to ozone stress is very much similar to pathogen stress and pathogen resistant crop varieties may also possess resistance to ozone stress.

**Fig 4:**
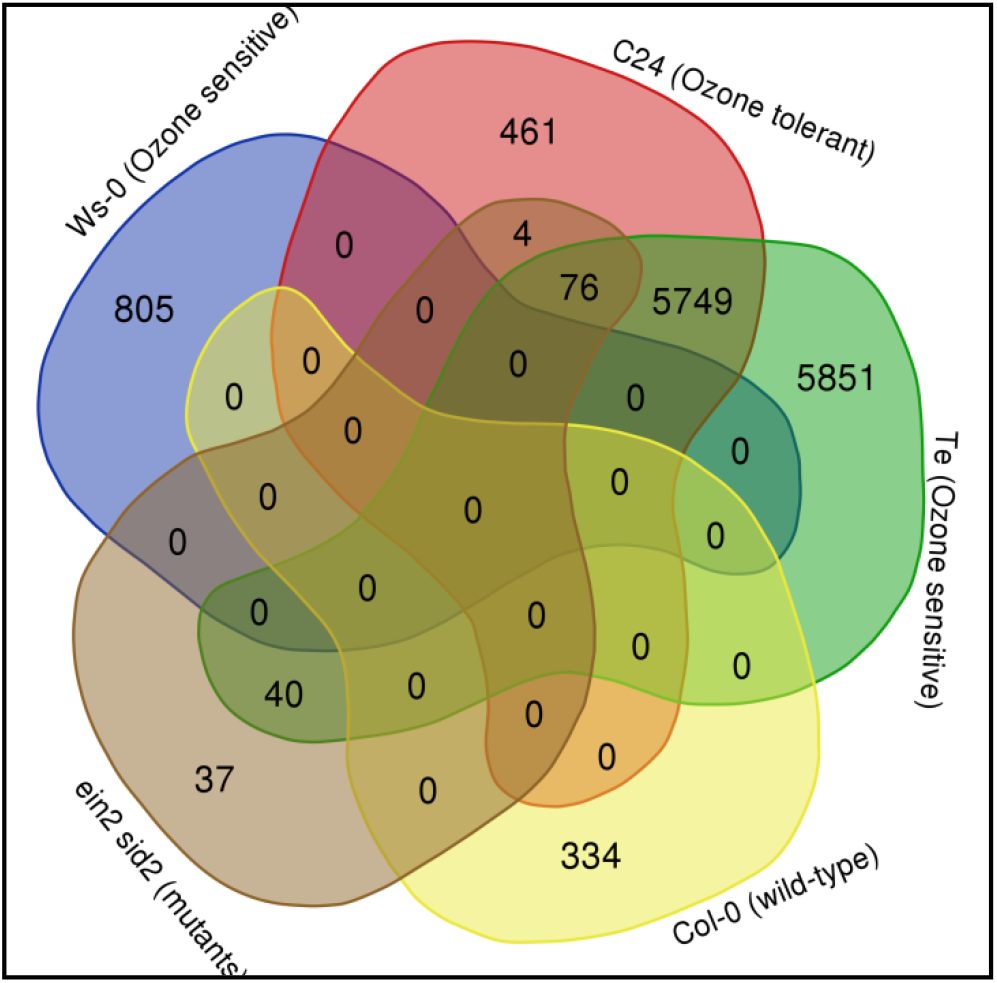
Venn diagram showing differentially expressed genes under ozone stress shared among representative ecotypes of *Arabidopsis thaliana*

A study on *A.thaliana* ecotype Col-0 and its mutants *ein-2* (ethylene signaling deficient) and *sid-2* (salicylic acid biosynthesis deficient) revealed that phytohormones play important role in protecting plants against ozone stress. The most responsive biological processes in wild type Col-0 were: i) removal of superoxide radicals, ii) cellular oxidant detoxification, iii) response to oxygen radicals, iv) L-ascorbate biosynthesis, v) icosanoid metabolism. The highest activity was observed for superoxide dismutase, oxido-reductase, catalase, and glutathione peroxidase in chloroplast and peroxisome (Fig S5) [Yoshida e al., 2009].

In most plant species the gene expression studies with respect to ozone stress have focused majorly on leaf tissue as leaves possess stomata through which ozone enter and elicit a rapid signaling response. But for the development of complete transcriptome map, analysis and identification of tissue- and organ-specific genes is quite important. In the case of ozone stress, gene expression studies were reported in *Glycine max L. Merr. cv.* 93B15 for specific tissues such as pods, flowers and seed coat. The analysis of ozone specific transcriptome data in leaves of ozone sensitive cultivar (Mandarin-Ottawa) and an ozone resistant cultivar (Fiskeby III) of *Glycine max* revealed difference in gene expression levels indicating variability in cultivars response to ozone stress [Whaley et al., 2015]. Another study on the leaves of ozone sensitive wild soybean accessions PI 407179 and PI 424007 and ozone resistant wild soybean accessions PI 424123 and PI 507656 indicated age dependent response of *Glycine max* to ozone stress. Older leaves exhibited reduction in transcript level of photosynthesis related genes whereas younger leaves majorly displayed the expression of defense related genes [Waldeck et al., 2017]. The gene expression study in flower and pod tissue of soybean revealed that both tissues have distinct transcriptome responses to ozone stress. Flower tissues exhibited the increased expression of genes encoding matrix metalloproteinases (MMPs) which are endopeptidases having role in programmed cell death and senescence. The pod tissue showed increased expression of xyloglucan endo-transglucosylase/hydrolase genes having possible role in increased pod dehiscence during ozone stress [Leisner et al., 2014]. However, the seed coat specific transcriptome of soybean under ozone stress revealed 148 differentially expressed genes in response to ozone stress indicating non significant transcriptional changes in seed coat [Leisner et al., 2017]. The tissue specific studies in *Glycine max L. Merr. cv. 93B15* revealed that different tissues in a species expressed unique set of genes under ozone stress. A very few number of genes were found overlapping between flower, pod and seed coat. A complete unique set of genes in the leaf tissue indicated distinct tissue specific patterns of gene regulation in the species (Fig 5).

**Fig 5:**
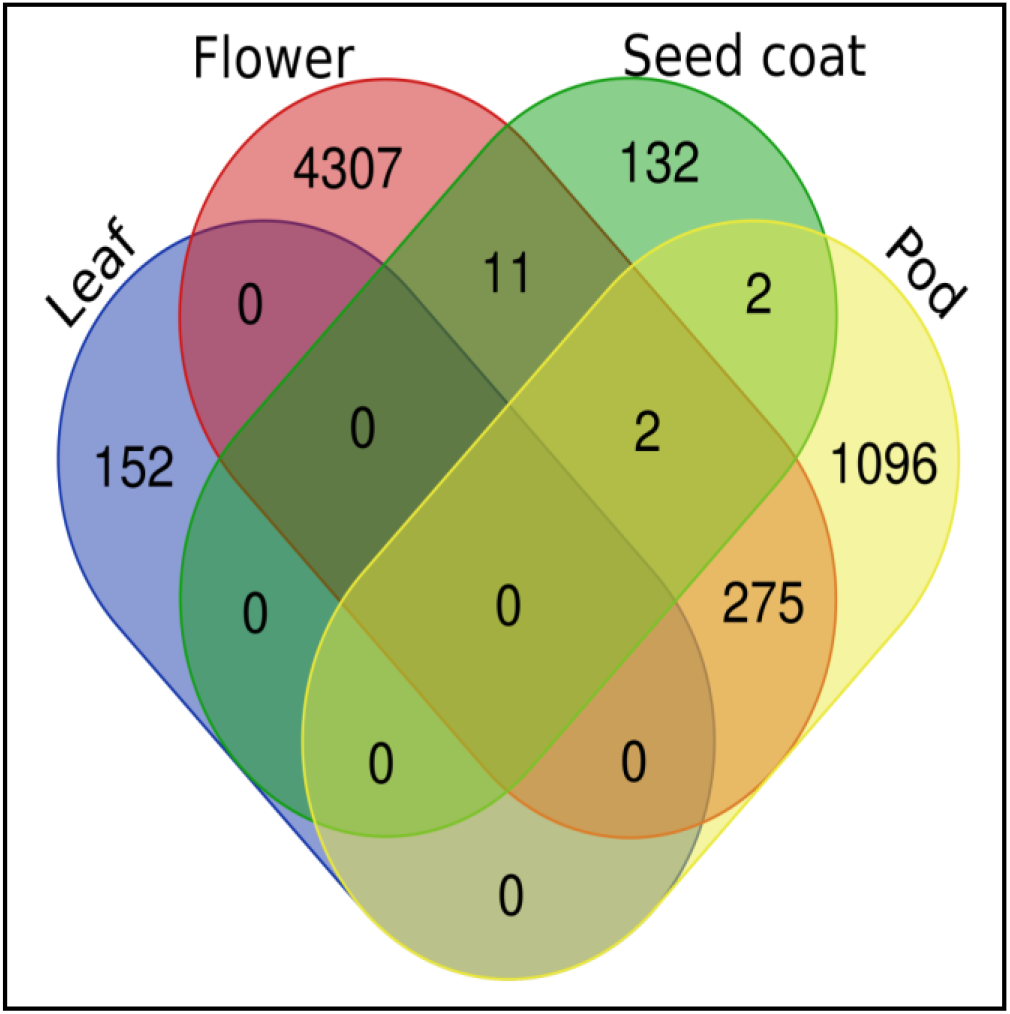
Venn diagram showing tissue specific unique and shared set of genes in *Glycine max L. Merr. cv. 93B15* under ozone stress

As evident from the available literature, the number of genes up-regulated and down-regulated under ozone stress is plant species-type and tissue-type specific and largely depends upon the cultivar chosen and dose and duration of the ozone stress. Therefore, different studies are required to be carried out for gene expression studies under different set of conditions to gain complete knowledge underlying the ozone stress response in plants.

Since the underlying ozone stress specific molecular mechanism through gene expression analysis have begin to unravel in recent years only, the gene expression data generated in different plant species is non-exhaustive and needs to be updated and expanded periodically. The current database is OSRGD ver. 1.1 having enormous scope to be updated in future. In future versions, the information of proteins encoded by the ozone stress responsive genes and micro-RNAs involved under ozone stress can be included.

## Conclusion and Future directions

Understanding of the gene expression in response to ozone stress in plants is important for the identification and development of ozone-tolerant varieties of important crops which can adapt and grow well in areas having frequent incidences of elevated ozone levels. Ozone Stress Responsive Gene Database (OSRGD ver. 1.1) is first such comprehensive and user-friendly reliable resource of ozone stress responsive genes which would aid the plant researchers and computational biologists working for genetic improvement of crops. The database will be updated regularly with the availability of new gene expression dataset and will be further improved in future with enhanced functionality to serve as a more valuable resource for facilitating research community working on crop breeding programmes.

## Acknowledgement

Authors thank Dr. Vivek Dhar Dwivedi, Scientist at Pathfinder Research and Training Foundation (PRTF), Greater Noida, Uttar Pradesh, India for his critical suggestions in database improvement. Authors are also thankful to University Grants Commission for fellowship to PY.

